# Advances in magnetic particle imaging: evaluating magnetic microspheres and optimized acquisition parameters for high sensitivity cell tracking

**DOI:** 10.1101/2024.12.21.629270

**Authors:** Samantha N. Flood, Paula J. Foster

## Abstract

**Motivation:** The sensitivity and resolution of magnetic particle imaging (MPI) depend on the choice of tracer and specific imaging parameters. For cell tracking applications with MPI, both the superparamagnetic iron oxide (SPIO) tracer and the cell labeling efficiency have a significant impact on MPI sensitivity and vary for different tracers.

**Objective:** This study compared three commercially available SPIO tracers (VivoTrax, Synomag-D and ProMag) and SPIO-labeled cells using magnetic particle relaxometry (MPR) and imaging. Further, the effect of imaging parameters (gradient field strength and drive field amplitude) on MPI signal strength, resolution, and cell detection limits, was evaluated.

**Results:** The peak MPI signal measured by MPR was much higher for Synomag-D compared to VivoTrax and ProMag. However, the signal for intracellular Synomag-D was significantly reduced. In contrast, the signal for ProMag, a micron-sized iron oxide (MPIO) particle, was not significantly different for free and intracellular particles. The cellular iron loading was higher for ProMag compared to Synomag-D. The total MPI signal measured from images of free and intracellular SPIOs was highest for ProMag. Varying imaging parameters confirmed that a lower gradient field strength and higher drive field amplitude improved tracer and cellular sensitivity.

**Conclusion:** These results show that the evaluation of tracers by MPR is not sufficient to predict the performance of all SPIO tracers; in particular, not for larger, polymer-encapsulated iron particles such as ProMag, or for SPIO tracers internalized in cells. Improvements in MPI sensitivity through lower gradient field strength and higher drive field amplitudes are associated with a trade-off in image resolution.

## Introduction

Cellular therapeutics are being used clinically for the treatment of several diseases and injuries. While cell therapy shows immense potential in preclinical studies there has been limited success for many clinical trials [1–3]. This is in part due to a lack of cell survival and retention post-administration with some studies showing that <10% of the therapeutic cells survive the first few hours after administration and <3% after 24 hours [4,5]. There remains many unanswered questions about why these therapeutic cells are dying and how to enhance their survival. Establishing a tool to track and assess therapeutic cells is essential to further the current state of cellular therapeutics and ideally transition more therapies into the clinic.

Cell tracking has been used to assess the administration, biodistribution, retention, and the potential efficacy of therapeutic cells *in vivo*. Historically, *in vivo* cell tracking has been done with imaging technologies such as bioluminescence imaging (BLI), magnetic resonance imaging (MRI), and nuclear imaging (positron emission tomography (PET) and single-photon computed tomography (SPECT)). Clinically, only MRI and nuclear medicine can be used *in vivo* [6]. Cell tracking with MRI requires the use of a contrast agent to label cells. The most commonly used agents are superparamagnetic iron oxide (SPIO) particles and perfluorocarbon nanoemulsions [7]. Iron-based MRI cell tracking suffers from medium sensitivity, low specificity and a lack of direct quantification. Fluorine-19 (^19^F) MRI is more specific and directly quantitative but has very low sensitivity with thousands of cells needed per voxel to acquire signal [8,9]. Nuclear imaging provides higher sensitivity than ^19^F MRI and is also directly quantitative, but the half-lives of the radioisotopes used make long-term tracking complex and deliver a low radiation dose to the patient and healthcare workers [6,10]. Evidently, there is a need for non-invasive, non-ionizing, sensitive, and directly quantitative imaging modality for cell tracking applications.

Magnetic particle imaging (MPI) is an emerging imaging modality that directly detects SPIO particles. One of the early applications of MPI has been *in vivo* cell tracking and has been performed in a variety of applications including the monitoring of cellular therapeutics, cell biodistribution, cell retention and clearance. More specifically, MPI cell tracking has been used to track stem/stromal cells [11–14], immune cells [15–20], cancer cells [21–24], and cellular components like extracellular vesicles [25,26]. The main advantage of using MPI for cell tracking, over other imaging modalities, is the ability to estimate cell number from images. MPI does not produce anatomical images but can be combined with other imaging modalities, such as CT or MRI, to provide spatial information. Currently, various groups are engineering human-size prototype MPI systems for clinical imaging [27–30]. It has been theorized, that an optimized clinical MPI device could detect as low as 25 pg of iron but with numerous hardware and safety limitations; it is, therefore, more practical to optimize sensitivity and resolution by optimizing tracer performance, cell labeling, and imaging parameters [31,32].

The physics of MPI are described in detail in the following comprehensive reviews: Knopp *et al.*, Neumann *et al.*, Irfan & Dogan, and Harvell-Smith *et al.* [33–36]. The MPI system consists of three main parts: a selection field, a drive field, and a receiver coil. The selection field is produced by two opposing magnets which produce a strong magnetic field gradient with a central region with a near-zero magnetic field, referred to as the field-free region (FFR). The drive field is an alternating magnetic field (AMF) that causes changes in the oscillation of SPIO particles producing the MPI signal. The signal is proportional to iron mass allowing for quantitative imaging [12,37]. Other advantages of this modality are high specificity, high sensitivity since there is no signal from biological tissues, and no limit to imaging depth since there is no signal attenuation from tissues.

The strength of the gradient field determines the inherent resolution of the MPI system; this is the resolution that arises only from the interaction between the gradient field and the SPIO particles. Spatial resolution can be improved by increasing the gradient field strength; doubling the gradient field strength will improve image resolution by a factor of 2 [38,39]. However, a higher gradient field strength reduces the scanning volume of the FFR, which reduces the MPI signal [40]. Lower drive field amplitudes have been shown to improve particle resolution but at a large cost in sensitivity. For example, Tay *et al.* showed that for the SPIO tracer VivoTrax, an 8-fold decrease in drive field amplitude is needed to improve resolution from 1.6 mm to 0.9 mm [41]. A few studies have evaluated or optimized imaging parameters for high resolution and sensitivity but only using simulations or experiments with samples of SPIO particles [38,41,42]. The effects of these parameters on imaging SPIO-labeled cells *in vitro* or *in vivo* have yet to be performed.

The formulation of SPIO tracers tailored for MPI has the potential to dramatically improve sensitivity and resolution. Numerous studies have characterized and compared the magnetic properties of commercial SPIO and MPI-tailored SPIO tracers [43–46]. The best MPI *in vivo* detection limits to date are 1000 cells (corresponding to 4.4 ng Fe) labeled with a commercially available SPIO tracer and 250 cells (corresponding to 7.8 ng Fe) with a custom-made SPIO tracer [45,47]. The MPI performance of SPIO particles depends on their magnetic properties, such as the saturation magnetization and the magnetic core size. The best performing SPIO tracers reported to date are monodisperse, single-core particles with core sizes of approximately 25 nm [48–50]. Larger magnetic microspheres, or micron-sized iron oxide (MPIO) particles, have magnetic properties and relaxation behaviour that is more complex [51,52]. Our lab has demonstrated the use of MPIOs for imaging cancer cells with MPI [22–24,53].

For cell tracking with MPI, the type of SPIO is also important for other reasons. First, it is essential that the SPIO particles effectively label the cells. Factors that affect the cellular uptake of SPIO particles include the surface coating and charge, hydrodynamic size, the cell type, cell size and phagocytic ability of the cell; all of which impact the total iron loading of cells [9,54–56]. Second, some SPIO tracers have more iron per particle. MPIO particles have been used for cell tracking by MRI to enable the detection of single cells *in vivo* and single particles in cells [57,58]. Increasing the intracellular iron content, with high uptake of SPIO particles or by using MPIO particles which have very high iron content, will improve MPI cellular sensitivity. Third, the performance of SPIO particles can be affected by their environment. The MPI signal for some SPIO tracers is reduced in more viscous environments and after cell internalization [47,55,56,59–62]. This occurs to different degrees with different SPIO tracers and cell types and is, therefore, an additional important consideration.

The main goals for this study were to assess MPIO particles for imaging cells with MPI, by comparing them to commercially available SPIO tracers, and to test the effects of changing individual MPI scanner-specific imaging parameters on image sensitivity, resolution, and cellular detection limits.

## Materials and Methods

### Superparamagnetic Iron Oxide (SPIO) Particles

Three commercially available SPIO particles were obtained: VivoTrax (Magnetic Insight Inc., Alameda, California, United States), Synomag-D (Micromod Partikeltechnologie, GmbH, Hamburg, Germany), and ProMag (Bangs Laboratories, Fishers, Indiana, United States). VivoTrax was chosen because it is the most commonly used SPIO for MPI cell tracking. It is a multicore particle with a bimodal size distribution of ∼30% 25-30 nm cores and ∼70% 5 nm cores, it has a carboxydextran coating and a reported hydrodynamic size of 62 nm. Synomag-D was chosen because it has previously been shown to have a higher MPI signal compared to VivoTrax, as measured by MP relaxometry (MPR) [43,47]. It is a multicore particle with a nanoflower substructure with a reported cluster core size of ∼30 nm, a dextran coating and a hydrodynamic size of 50 nm [46]. ProMag consists of multiple iron oxide cores encapsulated within an inert polymer matrix. The total diameter of each microsphere is reported to be 1 µm.

### MPI Relaxometry of Free SPIO Particles

MPR was performed using the RELAX^TM^ module on a Momentum^TM^ preclinical scanner (Magnetic Insight Inc., Alameda, California, United States). Relaxometry is a commonly used method to assess the magnetic properties of SPIO particles. The output of the RELAX^TM^ module is a point spread function (PSF) characteristic of the entire volume of a SPIO sample. The peak signal intensity of the PSF for each SPIO sample reflects the particle sensitivity and the full-width half maximum (FWHM) is related to the particle resolution. The FWHM in mT can be converted to mm by dividing by the gradient field strength (T/m). Free SPIO samples were made in triplicate for each tracer in 0.2 mL individual tubes (n = 3 for each SPIO tracer; ProMag = 8 µL, Synomag-D = 1.5 µL, VivoTrax = 12 µL). Different volumes were used to assess each SPIO tracer to not oversaturate the receive coil to the system since each tracer has a different iron concentration. To compare the PSFs for the different SPIO tracers, the signal was normalized by the iron concentration of the sample.

### Cell Preparation and Labeling

Human adipose-derived stromal cells (ASC) were cultured in a T75 cm^2^ flask at 37°C and 5% CO_2_ in Dulbecco’s modified Eagle’s medium F12 (DMEM F12) containing 10% fetal bovine serum (FBS) and 1% penicillin-streptomycin (PS) until 90% confluency. Cells were washed with phosphate buffer solution (PBS) to remove dead cells and residual media components immediately before cell labeling.

To label ASC with ProMag, 10 mL of complete medium and 54 µL of ProMag (4.79 µg Fe/µL) were added to culture at a final concentration of 25.9 µg/mL media. Following overnight incubation, cells were washed 3 times with PBS to remove unincorporated particles. To dissociate adherent cells, 5 mL of trypsin/disodium ethylenediaminetetraacetic acid (EDTA) was added, and the flask was placed in the incubator for 5 minutes. 5 mL of complete media was added to the dissociated cells then centrifuged at 300 g for 5 minutes. After centrifugation, the supernatant was aspirated then the cell pellet was resuspended in 10 mL PBS and centrifuged for 5 minutes. This washing and centrifugation in PBS was repeated 2 more times in effort to remove extracellular iron. Labeled ASCs were suspended in 2 mL PBS and incubated for 5 minutes in an EasySep™ magnet (Stemcell Technologies, Vancouver, CAN) to separate the iron-labeled cells from the unlabeled.

Cell counting was completed using the trypan-blue exclusion assay (Countess Automated Cell Counter; Invitrogen) to determine the concentration of cells in PBS. Cell samples were prepared in 0.5 mL Eppendorf tubes, containing 1, 2, 5, 10, 25, 50, 100, or 500 x 10^3^ cells pelleted by centrifugation.

To label ASC with Synomag-D, transfection agents were employed. Efficient cell labeling with Synomag-D requires the use of transfection agents [15,21,63]. Transfection agents, such as heparin and protamine sulfate, create complexes or aggregates with the SPIO particles through electrostatic interactions which promote endocytosis of the particles by cells [64]. In one tube, 2.5 mL of serum-free DMEM F12 and 60 µL of stock protamine sulfate (USP, 10 mg/mL) were added and vortexed. In another tube, 2.5 mL of serum-free DMEM F12, 90 µL of Synomag-D and 20 µL of stock heparin (1000 USP units/mL) were added and vortexed. The two tubes were combined and vortexed before the labeling mixture was added to the flask containing ASCs. After co-incubation for 4 hours, 5 mL of complete DMEM F12 was added to the flask and the cells were left to incubate overnight. The next day the cells were collected, and counted, and cell pellet samples made by the same method described for labeling with ProMag.

### Assessing Cell Labeling

Perls’ Prussian Blue (PPB) staining was performed to qualitatively assess iron labeling efficiency. Cell samples of 200,000 cells were added to glass slides using a Cytospin 4 centrifuge (Thermo Scientific) set at 1000 rpm for 5 minutes and then allowed to dry for 5 minutes. Next, the cells were placed in a fixative mixture of methanol and acetic acid (3:1) and left to sit for 5 minutes. The slides were rinsed in deionized water and then placed in a PPB solution. The PPB solution was made from 0.5 g PPB powder, 25 mL Milli-Q water, and 25 mL of 2% hydrochloric acid which was vortexed and then filtered into a glass slide holder. The slides were left in the PPB mixture for 30 minutes. PPB stains iron blue during histochemical staining because the ferric iron in the SPIO particles react with the potassium ferrocyanide in the stain to form ferric ferrocyanide which is a blue pigment. Another deionized water rinse was performed, and the slides were placed in a nuclear fast red stain that was created using 25 g aluminium sulfate hydrate, 500 mL Milli-Q water, and 0.5 g nuclear fast red powder which were mixed on a hot plate using a magnetic stir bar for about 1 hour. Nuclear fast red was used as a counter stain as it stains nuclear chromatin red and provides nonspecific background tissue and cell staining with pink. The slides were left in the nuclear fast red stain for 10 minutes before undergoing various steps of increasing concentrations of ethanol (70%, 95%, 100%) and then two xylene washes to dehydrate the sample. Using cytoseal protectant a glass slide coverslip was immediately placed after the last xylene wash. Slides were left to dry for 24 hours before microscope imaging was completed using the Echo 4 Revolve Microscope (California, USA).

### Imaging of Free SPIO Particles

Five samples of ProMag and Synomag-D were created in a dilution series. The ProMag samples include 9.58 µg, 4.79 µg, 2.40 µg, 1.20 µg, and 0.60 µg of iron. The Synomag-D samples include 10 µg, 5 µg, 2.5 µg, 1.75 µg, and 0.88 µg of iron. All samples were diluted in PBS in 0.2 mL individual tubes for a final volume of 3 µL. The samples were then taped to the imaging bed in the same location and imaged individually with the following parameters: 2D field of view (FOV) = 12 x 6 cm (z, x), 5.7 T/m gradient strength, 20 mT (x-channel) x 26 mT (z-channel) drive field amplitudes, 0° projection angle (coronal image), and imaging time of approximately 2 minutes per sample (standard scan mode on the Momentum^TM^ system). Multichannel imaging collects two sets of data that are averaged to form a 2D isotropic image. This data was used to evaluate the relationship between total MPI signal and iron mass for each SPIO tracer.

### Imaging of Labeled Cells

Triplicate samples of Synomag-D- and ProMag-labeled cells were prepared by adding 500,000 cells and PBS for a total volume of 150 µL and then pelleting by centrifugation in a 0.5 mL centrifuge tube. Cell samples were imaged individually using the same parameters as listed above for the free SPIO samples. These images were used to calculate the iron loading per cell and to normalize MPR of labeled cells. Next, six samples of Synomag-D- and ProMag-labeled cell pellets were prepared as described above in a dilution series (1,000,000, 500,000, 250,000, 100,000, 50,000, and 10,000 cells) and imaged with the same parameters. This data was used to evaluate the relationship between total MPI signal and iron mass for Synomag-D- and ProMag-labeled cells and for signal calibration.

### MPI Relaxometry of Labeled Cells

MPR was performed on the same triplicate 500,000 Synomag-D- and ProMag-labeled cell pellet samples that were imaged to study the effects of cellular internalization on the peak signal and resolution. To compare the peak MPI signals the PSFs were first normalized by the iron mass (determined from imaging). To compare particle resolutions the PSFs were normalized to the maximum peak signal value.

### *In Vitro* Testing of Imaging Parameters

Triplicate 2 µL samples of ProMag (9.58 ug Fe each) were prepared in 0.2 mL individual tubes. An advanced user interface was used which allowed for editing of pulse sequences on the Momentum^TM^ system. All samples were imaged individually using a 2D FOV = 12 x 6 cm. Certain imaging parameters were varied to determine the effect on the MPI signal. The gradient field strength was varied first. Images were acquired with the standard isotropic imaging mode (used for all imaging described above) which used a 5.7 T/m gradient field strength and 20 mT (x-channel) x 26 mT (z-channel) drive field amplitudes and then with a 3.0 T/m gradient field strength and 20 mT (x-channel) x 26 mT (z-channel) drive field amplitudes. The drive field amplitudes were varied next. Images were acquired with a gradient field strength of 5.7 T/m and drive field amplitudes of either 5 mT (x-channel) and 5 mT (z-channel) or 20 mT (x-channel) and 20 mT (z-channel) mT. The same imaging was performed again using ProMag-labeled cell pellets containing 100,0000, 50,000, 25,000, 10,000, 5,000, 2,000, and 1,000 cells, prepared as described previously.

### Flow Cytometry

Before *in vivo* MPI the viability of cells labeled with ProMag was evaluated by flow cytometry using the SYTOX^TM^ Blue dead cell stain (ThermoFisher Scientific, Mississauga, Ontario, Canada). SYTOX^TM^ Blue is a high-affinity fluorescent, cell impermeant nucleic acid stain that selectively binds DNA in dead cells with compromised plasma membranes. ProMag-labeled and unlabeled ASC were stained with 1 µL of SYTOX^TM^ Blue in 300 µL staining buffer (PBS with 2% FBS). A heat-shock control was used as a positive control for SYTOX^TM^ Blue staining. The positive control was created by removing 25% of the cells in the sample tube and heating the removed cells in a water bath at 56°C for 10 minutes and then placing them on ice for 1 minute before adding them back into the sample tube for staining. A CytoFLEX S V4-B2-Y4-R3 Flow Cytometer (Part Number C09766; Beckman Coulter Canada LP, Mississauga, Ontario, Canada) was used to detect SYTOX^TM^ Blue staining (80 mW 405 nm Violet laser excitation, emission in 450/45 nm filter).

### *In Vivo* Testing of Imaging Parameters

Female nude (NU/NU) mice (Charles River, Canada or USA) were obtained and cared for in accordance with the Guidelines for Care and Use of Laboratory Animals of Western University Canada and animal procedures and experiments were approved by the Animal Use Ethics Committee of Western University’s Council on Animal Care. Prior to imaging, mice were fasted for 12 hours with only water, corn bedding, and a laxative in their cage to reduce gastrointestinal iron signal in images which can interfere with the detection of low MPI signal [23]. Mice were anesthetized with 2% isoflurane gas in oxygen throughout injections and imaging. Cell samples of 5,000, or 2,000 ProMag-labeled ASC, suspended in 50 µL of PBS, were injected subcutaneously into the back of the mice (n = 2 for 5,000, n = 2 for 2,000 cells). *In vivo* 2D MPI was performed immediately after cell injection using the optimal values for gradient field strength and drive field amplitude determined from the *in vitro* cell sample experiments (3.0 T/m gradient field strength and 20 mT x 20 mT drive field amplitudes) with the addition of 3D imaging (35 projections).

### Statistical Analysis

Excel (version 2409) was used to analyze MPR to assess the strength of the linear relationship between the known iron mass or cell number and the total MPI signal using the Pearson correlation. Statistical analyses for MPR and free SPIO sample parameter testing were performed using GraphPad Prism (version 9). A one-way analysis of variance (ANOVA) and Tukey’s honestly significant difference (HSD) test for multiple comparisons were performed to compare the maximum signal and FWHM (resolution) measured for free and intracellular SPIO by MPR. A p-value of 0.05 was used to determine statistical significance unless otherwise indicated. An unpaired t-test was used to analyze differences between the MPI signal for free SPIO sample parameter testing. Line profiles were generated using MagImage software (Magnetic Insight, Inc.) and analyzed using Excel (version 2409).

### Image Analysis

MPI data sets were visualized and analyzed using Horos imaging software. Horos is a free and open-source code software program that is distributed free of charge under the LGPL license at https://Horosproject.org and sponsored by Nimble Co LLC d/b/a Purview in Annapolis, MD USA. 2D images of the empty sample bed were acquired for every set of unique imaging parameters. The standard deviation of background noise (SD noise) was measured from a region of interest (ROI) drawn over the entire FOV. To quantify signal from images of free SPIO samples and labeled cells a threshold of 5 times the average SD noise was used to mask the lower amplitude signal and yield a reliable measurement of MPI signal [37]. This imaging criterion is based on MPI signal with a signal-to-noise ratio (SNR) >5 according to the Rose Criterion [65,66]. The total MPI signal was calculated as the mean MPI signal multiplied by the delineated ROI area. Linear regression was performed for MPI calibration (known iron mass vs. measured MPI signal) to determine the calibration equation. This line is forced through the origin, under the assumption that background MPI signal, without a sample of iron, has an average of 0. Pearson’s correlation was conducted for MPI (known iron mass or number of cells vs. measured MPI signal). The iron mass was calculated from images by dividing the total MPI signal by the slope of the calibration line. Cell detection limits were defined as the minimum number of ASC detected with SNR > 5. Thus, cells with signal below the 5*SD noise criteria were considered undetected.

## Results

### Comparing ProMag with Synomag-D and VivoTrax

MPR shows Synomag-D produces superior sensitivity and resolution compared to ProMag and VivoTrax. The maximum signal from Synomag-D was 2.7 times higher than ProMag and 4.4 times higher than VivoTrax (Figure 1A). The measured FWHM for Synomag-D was 8.9 mT compared to 14.7 mT for VivoTrax and 24.3 mT for ProMag (Figure 1B). For a gradient strength of 5.7 T/m, this translates to a spatial resolution of 1.56 mm for Synomag-D, 2.58 mm for VivoTrax and 4.26 mm for ProMag. Further experiments with labeled cells evaluated only Synomag-D and ProMag.

**Figure 1:**
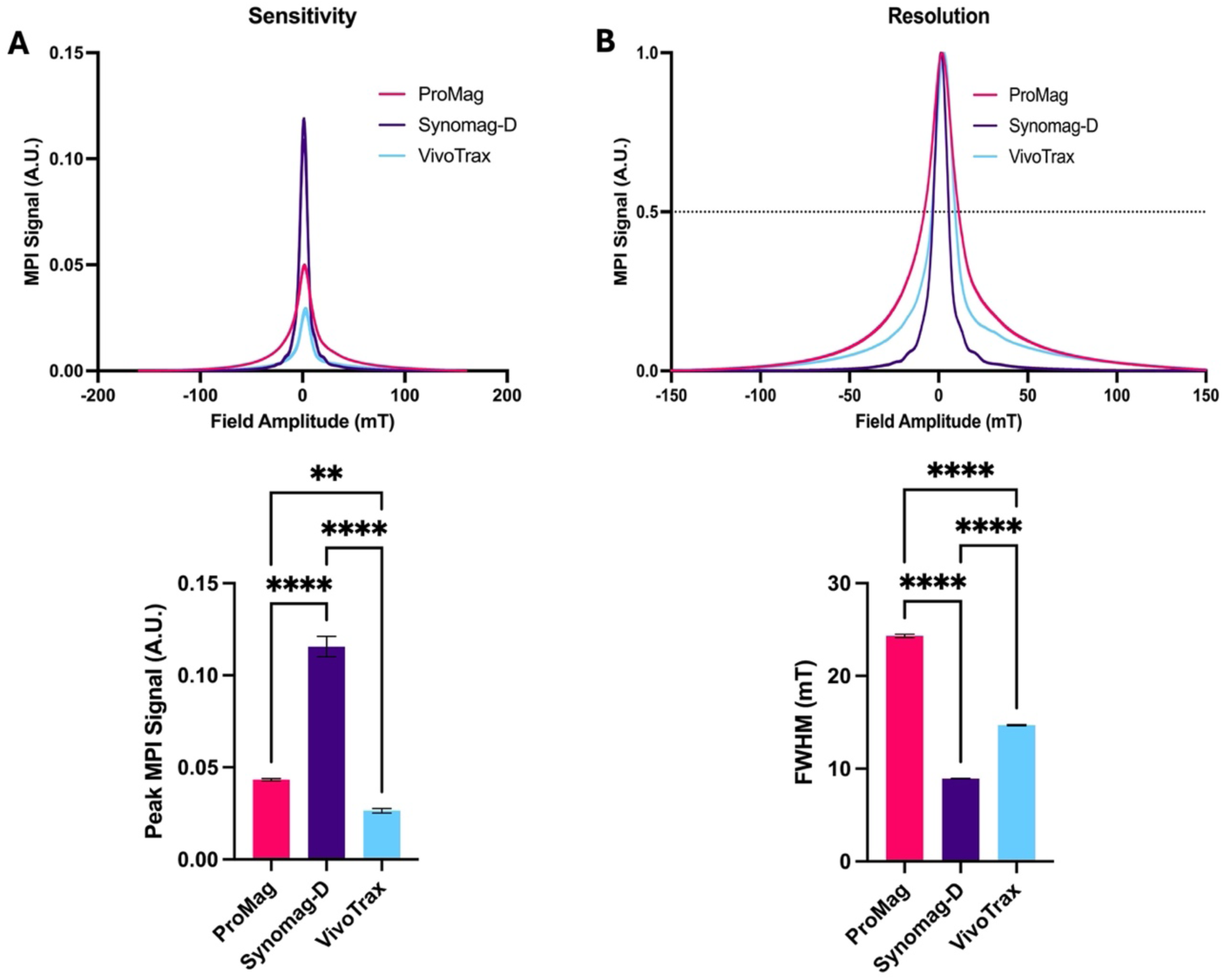
MPR showing the sensitivity and resolution for free Synomag-D, ProMag and VivoTrax. A) PSF and bar plot showing peak signal intensity normalized to iron mass (sensitivity) of 3 free SPIO tracers. Free Synomag-D had higher MPI signal compared to other SPIO tracers tested. B) PSF and bar plot showing FWHM (resolution) of 3 free SPIO tracers. Free Synomag-D had a smaller FWHM and thus better resolution than other SPIO tracers tested (Ordinary one-way ANOVA, F_2,6_ = 620.2, P < 0.0001; Tukey HSD, ns – p >0.05, ** - 0.0019, **** - p ≤ 0.0001).

### Cell Labeling Efficiency with ProMag and Synomag-D

PPB staining confirmed ASC labeling with ProMag and Synomag-D (Figure 2).

**Figure 2:**
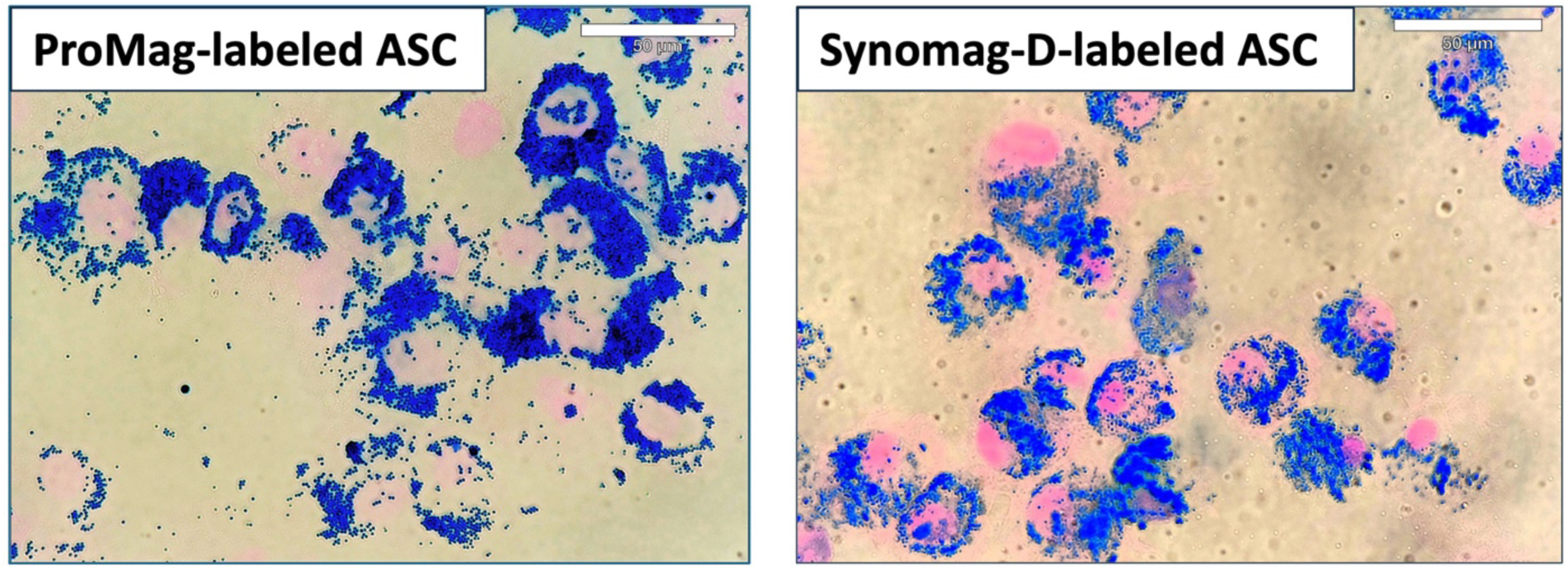
PPB staining of ProMag- or Synomag-D-labeled human ASCs where iron is stained blue and a counterstain of nuclear fast red stains the nucleus of cells pink.

MPI signal was strongly correlated with iron mass for both Synomag-D (R^2^ = 0.985) and ProMag (R^2^ = 0.998) (Figure 3A). The slope of the line for free ProMag was 1.72 times higher than free Synomag-D indicating higher sensitivity. The iron loading per cell measured from triplicate samples of 500,000 Synomag-D- and ProMag-labeled cells was 12 pg/cell for Synomag-D and 22 pg/cell for ProMag. The total MPI signal measured from images of cell samples was strongly correlated with cell number for both Synomag-D (R^2^ = 0.986, slope = 0.0002 A.U./cell) and ProMag (R^2^ = 0.994, slope = 0.0006 A.U./cell) (Figure 3B). The slope of the line for ProMag-labeled cells was higher than Synomag-D indicating higher sensitivity.

**Figure 3:**
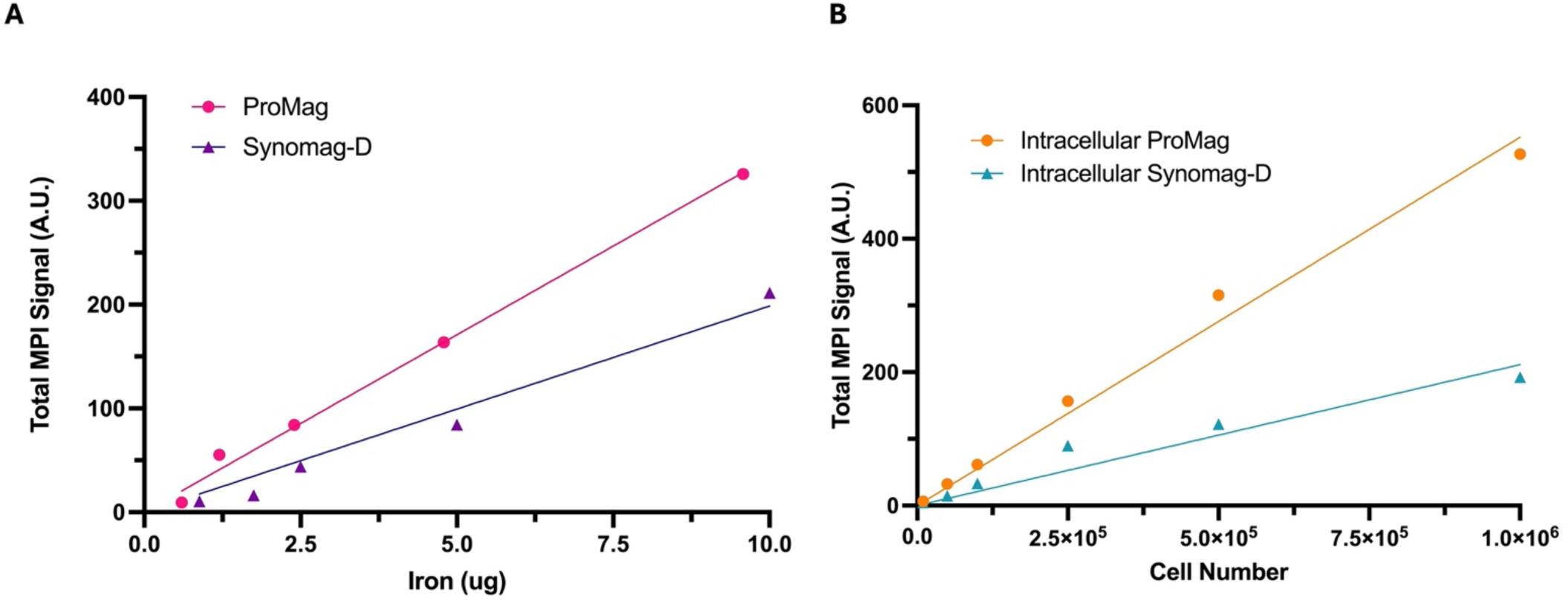
Total MPI signal measured from images versus known iron mass and cell number. The known iron mass and cell number are strongly correlated with the total MPI signal for both SPIO tracers. A) The slope of the line for free ProMag is steeper than free Synomag-D indicating higher particle sensitivity. B) The slope of the line for intracellular ProMag is steeper than intracellular Synomag-D indicating higher sensitivity.

Figure 4 shows MPR results for Synomag-D- and ProMag-labeled cells, compared to MPR of the free SPIO samples. The peak signal for Synomag-D decreased by ∼60% when in cells (Figure 4A). A reduction in the MPI signal after cell labeling, when SPIO particles aggregate, degrade or are immobilized, has been reported previously [60,67,68]. Importantly, there was no significant difference in the peak signal for free ProMag versus cells labeled with ProMag. Ultimately, there was no significant difference in the peak signal for cells labeled with Synomag-D or ProMag.

**Figure 4:**
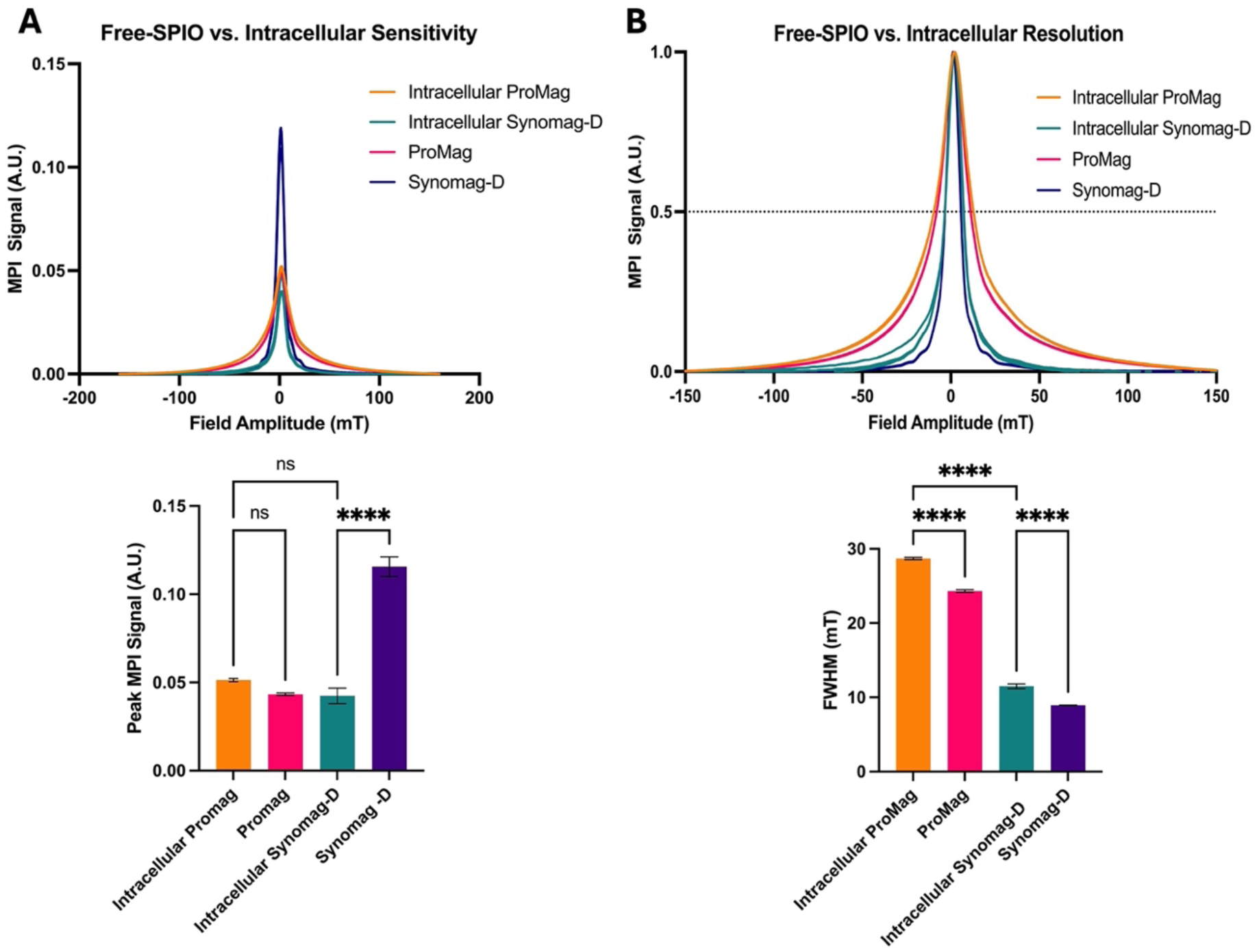
MPR data for free Synomag-D, ProMag, and ASCs labeled with each SPIO tracer. A) PSFs and bar plot showing peak signal intensity normalized to iron mass (sensitivity). ProMag signal did not change significantly once internalized into cells while Synomag-D signal dropped significantly. Once internalized, the signals from ProMag and Synomag-D were not significantly different from one another. B) PSFs and bar plot showing FWHM (resolution) of free SPIO samples and labeled cells. Resolution is worse for intracellular SPIO samples. Free Synomag-D and intracellular Synomag-D have a smaller FWHM and thus better resolution than free ProMag and intracellular ProMag respectively (Ordinary one-way ANOVA, F_3,8_ = 289.7, P < 0.0001; Tukey HSD, ns – p >0.05, **** - p ≤ 0.0001).

The resolution decreased for both SPIO tracers when internalized in cells. The FWHM for cells labeled with Synomag-D was 11.5 mT compared to 28.7 mT for ProMag (Figure 4B). For the highest gradient strength on the Momentum^TM^ system (5.7 T/m) this translates to 2 mm for Synomag-D and 5 mm for ProMag.

### Optimizing MPI Parameters for Cellular Sensitivity and Resolution

Experiments to determine how modifying imaging parameters affected the MPI signal and resolution were performed only with ProMag. This was based on the results of MPR and because of efficient cell labeling without the use of transfection agents.

The total MPI signal measured from images of free ProMag samples acquired with high and low gradient field strengths is shown in Figure 5A. The average total MPI signal was 2.3 times higher for images acquired with a gradient field strength of 3.0 T/m compared to 5.7 T/m. The increase in sensitivity achieved with a lower gradient strength comes with a trade-off in resolution. This can be appreciated in the representative images and line profiles in Figure 5B. The image resolution measured from the line profiles was 7.0 mm for the images acquired with a 3.0 T/m gradient strength and 3.7 mm for 5.7 T/m, which agrees with the expected linear scaling of resolution with gradient strength.

**Figure 5:**
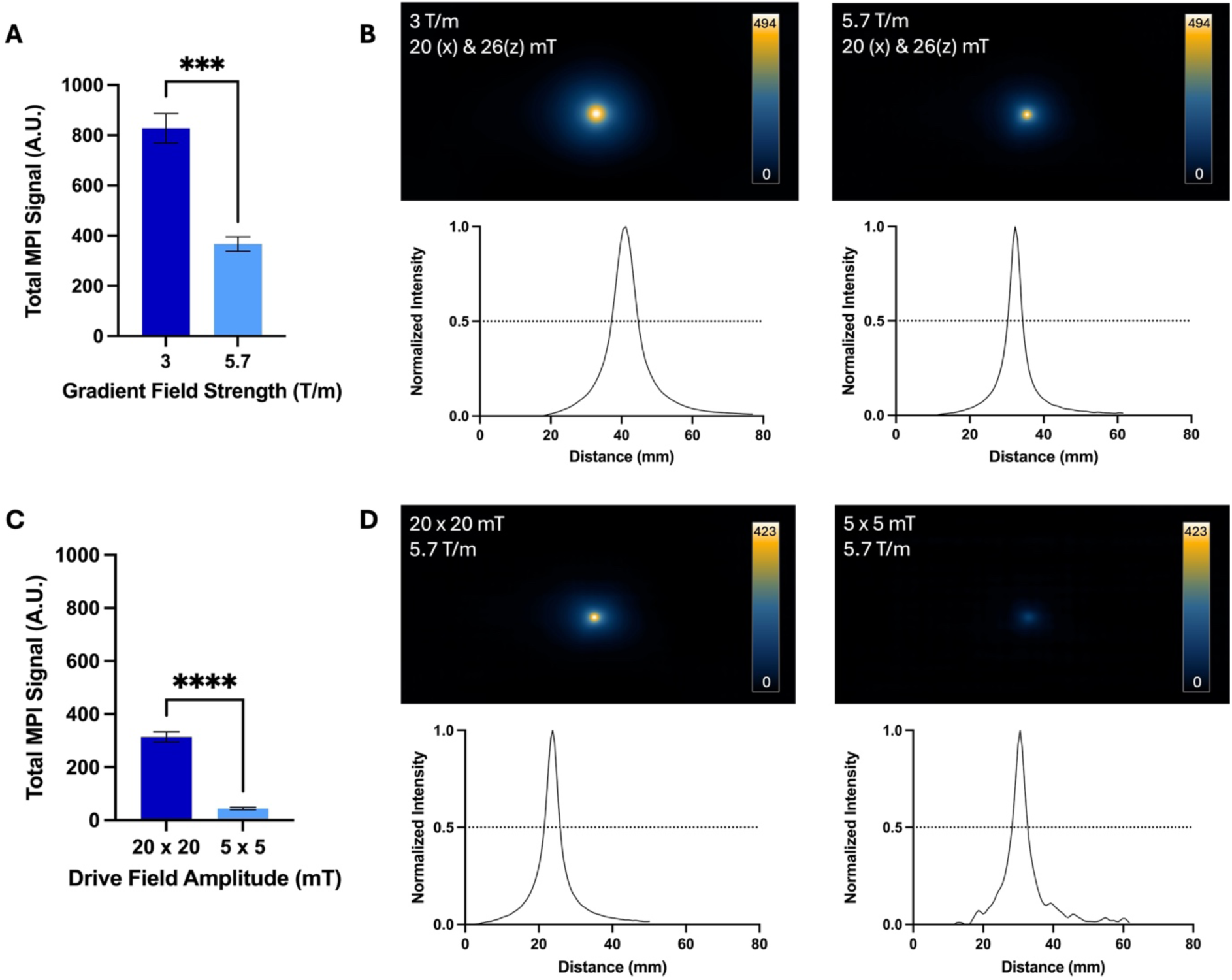
Comparison of images acquired with high and low gradient field strengths and high and low drive field amplitudes for free ProMag samples. A) The average total MPI signal was 2.3 times higher for images with the 3.0 T/m gradient field strength compared to 5.7 T/m. B) Representative 2D projection images are displayed with the scale set to the maximum signal value of the image acquired with 3.0 T/m. The lower resolution of images acquired with 3.0 T/m can be appreciated by the larger signal area and wider line profile compared to images acquired with 5.7 T/m. C) The average total MPI signal was 7 times higher for images acquired with the high drive field amplitude (20 x 20 mT) compared to the lower drive field amplitude (5 x 5 mT). D) Representative 2D projection images are displayed with the scale set to the maximum signal value of the image acquired with 20 x 20 mT. The signal in images acquired with 20 x 20 mT drive fields occupied a larger area compared to images acquired with 5 x 5 mT, indicating lower resolution.

The total MPI signal measured from images of free ProMag samples acquired with high and low drive field amplitudes is shown in Figure 5C. The average total MPI signal was 7 times higher for images acquired with drive field amplitudes of 20 x 20 mT compared to 5 x 5 mT. Again, with the increase in signal strength there is a trade-off in resolution which can be seen in representative images in Figure 5D. The image resolution measured from the line profiles was 4.0 mm for the images acquired with 20 x 20 mT drive field amplitudes and 3.7 mm for 5 x 5 mT.

The images and total MPI signal for ProMag-labeled ASCs acquired with high and low gradient field strengths are shown in Figure 6. As few as 1000 cells could be detected and quantified from 2D images acquired with a 3.0 T/m gradient field strength (the lowest number imaged). With the higher gradient field strength of 5.7 T/m as few as 5,000 cells could be detected. For cell numbers which could be detected with both gradient field strengths the average total MPI signal was 2.3 times higher for images of cells acquired with a gradient field strength of 3.0 T/m compared to 5.7 T/m. This result agrees with that observed for the free ProMag samples.

**Figure 6:**
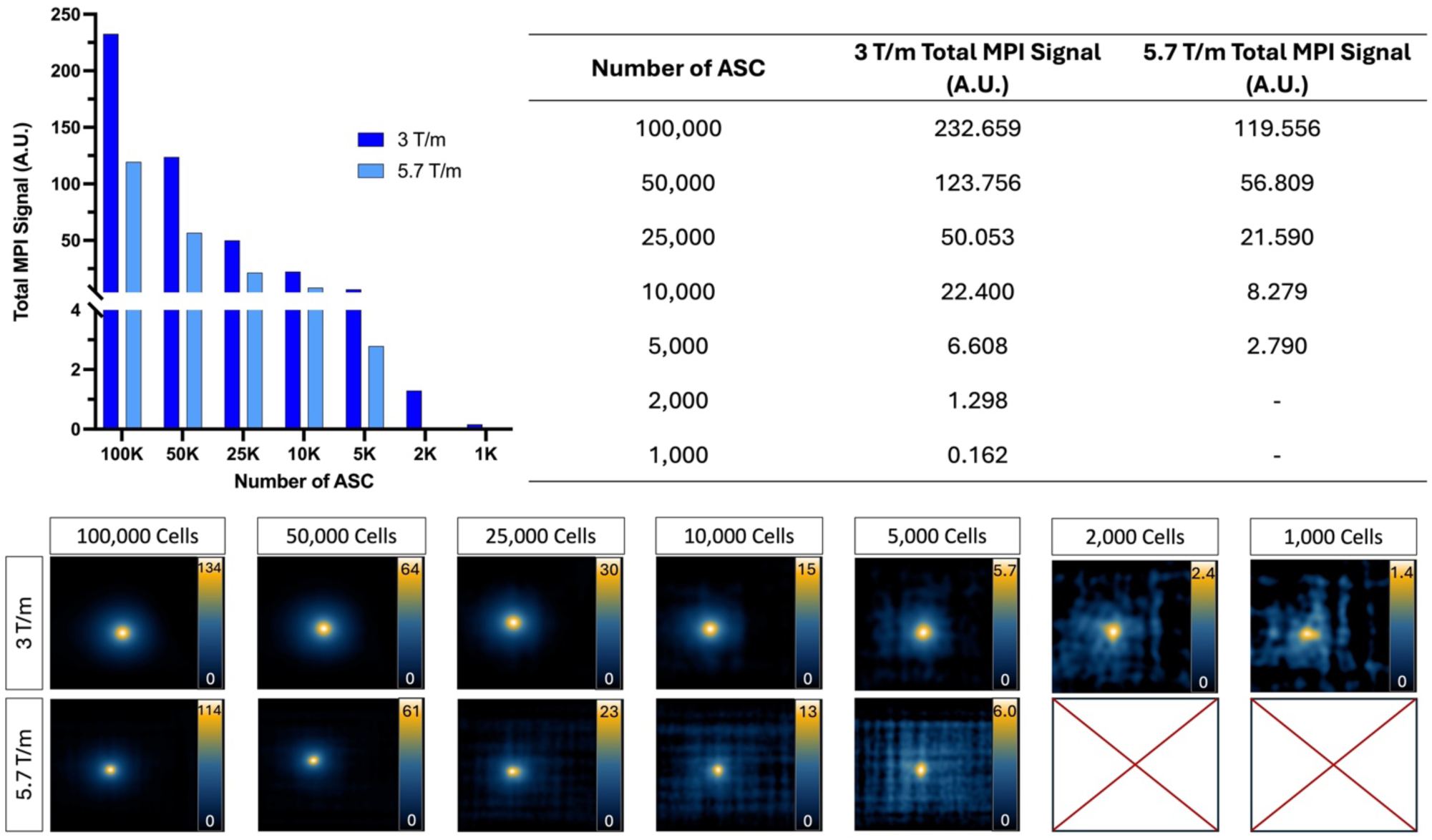
2D MPI of ProMag-labeled ASC pellets imaged using a low (3.0 T/m) or high (5.7 T/m) gradient field strength (drive field held constant at 20 x 26 mT). When using a high gradient field strength, as few as 5,000 cells could be detected. In comparison, when using a low gradient field strength as few as 1,000 cells could be detected showing improved sensitivity. Scales are set from zero to the maximum signal value.

The images and total MPI signal for ProMag-labeled ASCs acquired with high and low drive field amplitudes are shown in Figure 7. With the higher drive field strength of 20 x 20 mT as few as 5,000 cells could be detected. With the lower drive field amplitude of 5 x 5 mT the lowest cell number detected was 50,000 cells. The total signal calculated for the 100,000 and 50,000 cell samples was 14 and 24 times higher for images acquired with the 20 mT drive fields compared to 5 mT.

**Figure 7:**
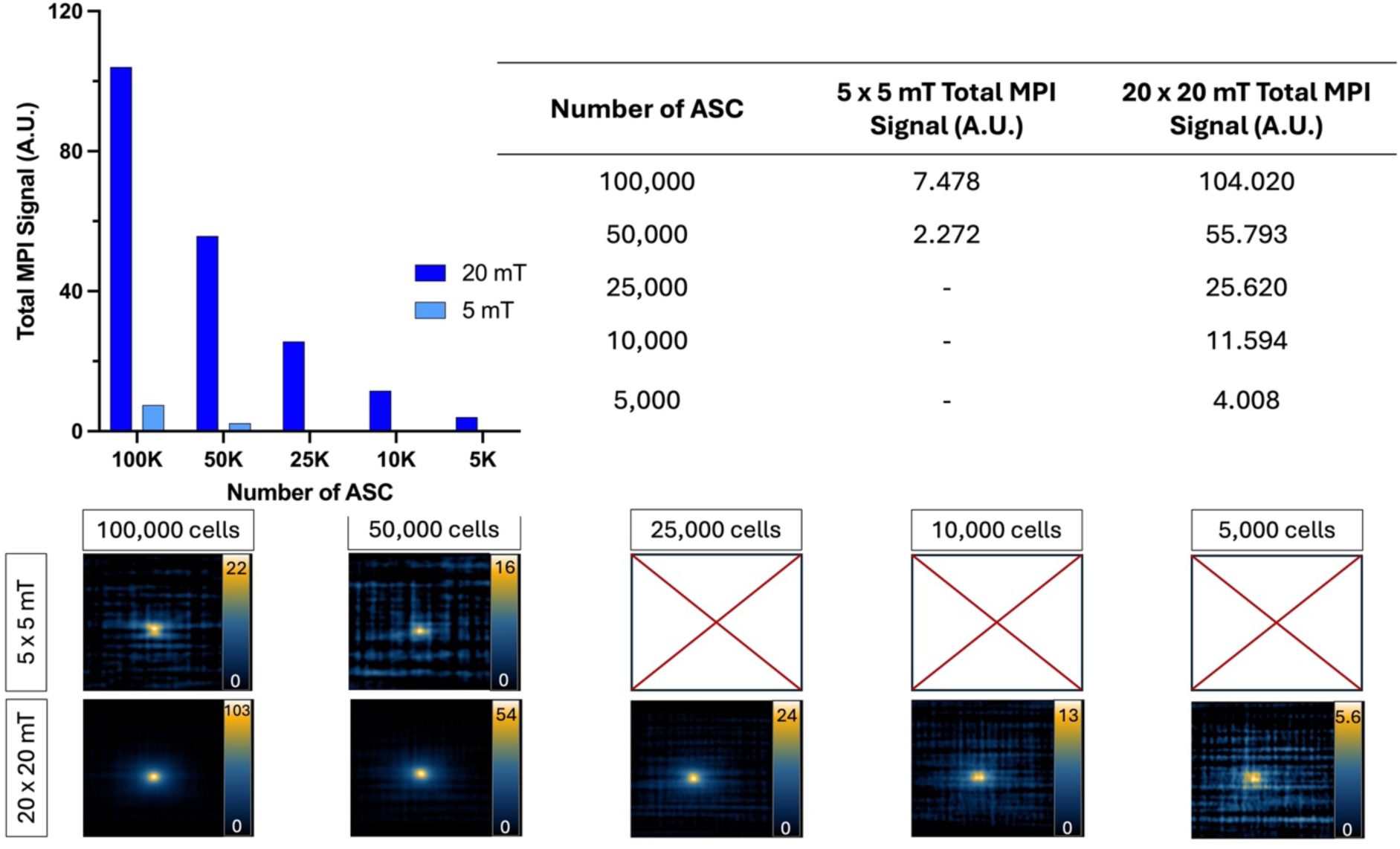
2D MPI detection of ProMag-labeled ASC pellets using a low (5 x 5 mT) or high (20 x 20 mT) drive field amplitude (gradient field strength constant at 5.7 T/m). When using a low drive field amplitude, as few as 50,000 cells could be detected. In comparison, when using a high drive field amplitude as few as 5,000 cells could be detected showing improved sensitivity. Scales are set from zero to the maximum signal value in each image.

### Viability of ProMag-labeled Cells

Flow cytometry was used to compare the viability of unlabeled and ProMag-labeled ASCs (Figure 8). The Sytox Blue viability assay showed a small decrease in viability for ProMag-labeled cells; there was a difference in viability of 15% between unlabeled (Figure 8A) and labeled cells (Figure 8B).

**Figure 8:**
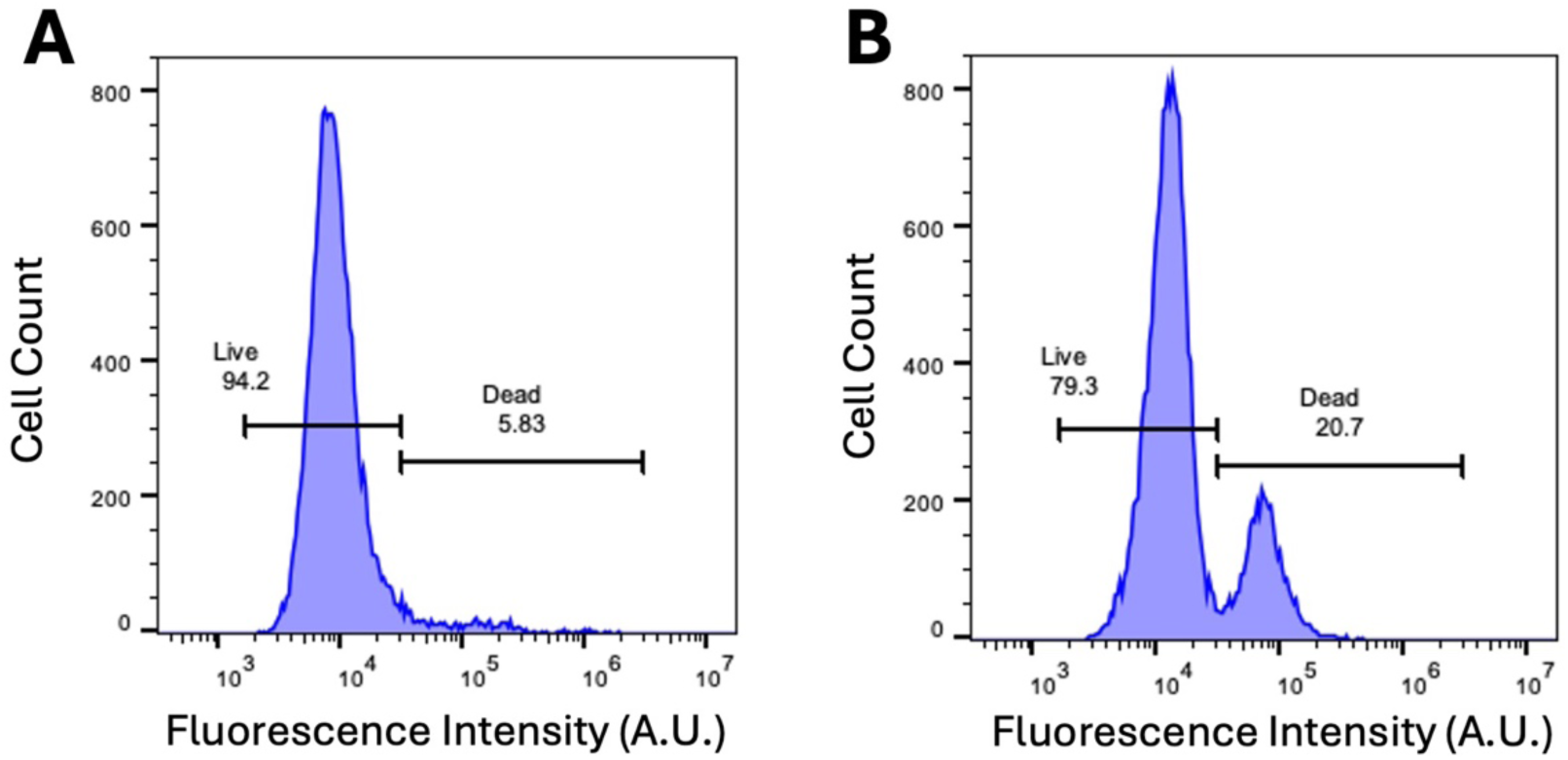
Sytox Blue dead cell stain. A) Flow cytometry of unlabeled ASCs with Sytox Blue viability assay dye reveals cell viability to be 95%. B) Flow cytometry with Sytox Blue viability assay after labeling ASCs with ProMag reveals a viability of 80%.

### *In Vivo* Testing of Imaging Parameters

Mice were imaged using the gradient field strength and drive field amplitude which gave the highest sensitivity in vitro experiments; 3.0 T/m and 20 x 20 mT (Figures 6 and 7). No signal was detectable in 2D MPI of mice injected with 2,000 or 5,000 cells. Figure 9A shows a representative 2D image of a mouse which was injected with 5,000 ProMag-labeled cells. MPI signal can be seen in the gastrointestinal region (arrow). This unwanted signal caused by iron in mouse feed has been observed before and can shadow regions of interest with low signal due to limited dynamic range [23]. However, when the same mouse was imaged with 3D MPI (35 projections) signal could be detected from the 5,000 ProMag-labeled cells at the ROI (arrow, Figure 9B). No signal was detectable with 3D MPI of mice injected with 2,000 cells.

**Figure 9:**
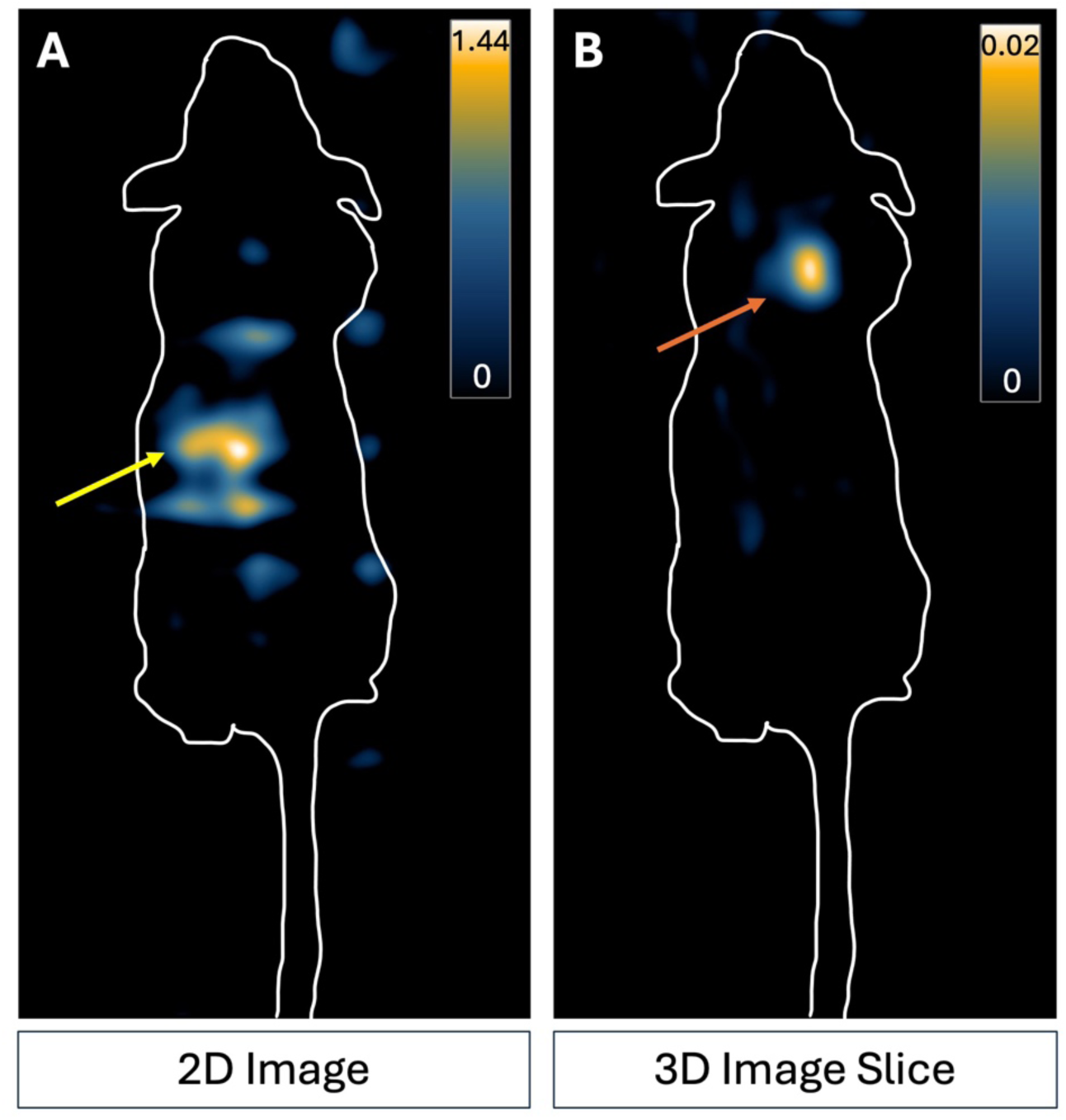
MPI images of a representative nude mouse which received a subcutaneous injection of 5,000 ProMag-labeled ASCs to the back of the neck. A) 2D imaging did not allow distinct visualization of the cell signal in comparison to the background signal. Signal present is due to iron in mouse feed in the gut (yellow arrow). B) The cells could be detected with 3D MPI. An increase in averaging (3D imaging, 35 projections) made the cell sample (indicated by the orange arrow) visible. Scale is the full dynamic range of the 2D image and 3D image slice.

## Discussion

### SPIO Tracer Comparison for High Sensitivity in MPI

In this study, the performance of SPIO tracers were evaluated by MPR and MPI. When comparing particles using MPR, Synomag-D had the highest peak signal. This agrees with several other studies that have shown that Synomag-D outperforms other commercial or newly synthesized SPIO tracers [43,46,47]. However, the peak signal for intracellular Synomag-D decreased by 60%. Previous studies have shown that the magnetic behaviour of SPIO particles is strongly influenced by the local environment. This can be due to changes in viscosity or aggregation of particles after cellular internalization, both of which are known to increase particle interactions and slow Brownian relaxation leading to decreased resolution and reduced signal [55,59,61,69]. The use of transfection agents to label cells, which was necessary for Synomag-D, has also been shown to reduce the MPI signal [47,55,63]. Gevaert *et al.* showed that the peak signal measured by MPR was lower for Synomag-D mixed with heparin and protamine sulfate compared to Synomag-D alone and was further reduced when the mixtures were internalized into dendritic cells [47]. This cell labeling method creates complexes of iron particles and transfection agents which increases the hydrodynamic size of particles and inhibits the signal. Notably, the peak signal measured by MPR for free ProMag was not different from ProMag-labeled cells. This may be because the encapsulation of the iron limits particle interactions and aggregation of ProMag in cells. Ultimately, the peak MPR signal was not different for intracellular Synomag-D and intracellular ProMag.

When comparing particles with MPI, ProMag samples had higher total MPI signal (per unit mass) compared to Synomag-D samples. This was not expected based on the peak signals measured by MPR. Guzy *et al.* observed something similar when comparing MPR and MPI of small versus large particles [67]. They hypothesized that the reduced peak signal measured by MPR and increased total signal measured by MPI for large particles is due to the decreased magnetization response. A larger magnetic field change is needed to saturate the magnetization of MPIO particles, and this is only achieved for MPI where the particles experience strong fields as the FFR is moved over the ROI. This data shows that the extrapolation of MPR results to MPI is not always straightforward, especially for large particles. For MPI of labeled cells, the total signal was again higher for ProMag compared to Synomag-D, however, the differences were greater than for the free SPIO samples. This is likely because here the total signal was plotted against cell number and the ProMag cell samples contain more iron than the Synomag-D cell samples.

These findings along with other benefits of using polymer-encapsulated MPIO tracers indicate that magnetic microspheres should be investigated further for their value as MPI tracers. The other important benefits of MPIO tracers for preclinical cell tracking include the fact that many cell types can be easily labeled with MPIO particles without the need for transfection agents with high cellular iron loadings (10-100 pg/cell) [57,58,70–73] and the potential for longitudinal imaging with MPIOs which are not rapidly degraded in cells [67]. While commercially available MPIOs such as ProMag are not biocompatible or biodegradable and, therefore, not suitable for clinical MPI, there has been progress in the synthesis of biocompatible MPIO [74,75]. Furthermore, the U.S. Food and Drug Administration has a minimum acceptable cell viability of 70% required for somatic cellular therapy new drug applications and our finding that 80% of cells are viable after labeling ASC with ProMag meets this clinical cutoff [76].

### Testing Imaging Parameters for Optimal Sensitivity and Improved Cellular Detection Limits

The effects of varying the gradient field strength and the drive field amplitude on the total signal and image resolution were evaluated by MPI for free ProMag samples and ProMag-labeled cell samples. Previous studies have tested changes in these parameters for other SPIO tracers in solution but to the best of our knowledge, there have not been experiments performed with cells or *in vivo*. Increasing the gradient field strength is expected to reduce the MPI signal and improve spatial resolution in a linear fashion. Our results for free ProMag and ProMag-labeled cells followed this prediction. The total MPI signal was approximately double when the gradient field strength was halved. The measurement of total MPI signal is important for quantification. Total signal is calculated by multiplying the mean MPI signal in the ROI by the ROI area and a calibration can then be used to convert total MPI signal to iron content. The peak MPI signal and the mean MPI signal were similar for images of free ProMag and ProMag-labeled cells acquired with gradient field strengths of 3.0 T/m and 5.7 T/m. However, the area was 2 times larger for images acquired with 3.0 T/m which resulted in the total MPI signal also being 2 times higher. The larger area is related to the lower spatial resolution at 3.0 T/m. For the Momentum^TM^ MPI scanner which uses a field free line (FFL) a lower gradient field strength results in a wider FFR which lowers resolution but collects more signal. As few as 1,000 cells (the lowest number imaged) could be detected with the lower gradient strength and 5,000 cells could be detected with the higher gradient strength.

Optimization of the drive field can also improve MPI signal and resolution. MPI theory and previous work have shown that the peak signal strength is directly proportional to the drive field amplitude, but only when neglecting the effects on relaxation. Most studies that have focused on the drive field are concerned with adjusting the imaging sequence to minimize relaxation effects for improved resolution. The results of such studies show that low drive field amplitudes improve resolution but at a significant cost of decreased sensitivity [38,41]. The effect of the drive field amplitude on large polymer-encapsulated iron oxide particles has not been evaluated before. We compared images acquired with drive field amplitudes of 5 x 5 mT and 20 x 20 mT. The mean signal was similar for both conditions and the area was approximately 7 times larger for the higher drive field amplitude resulting in the total MPI signal being 7 times higher. The image resolution was not significantly different for the two different drive field amplitudes. We hypothesize that the encapsulation of iron particles in ProMag suppresses Brownian relaxation limiting the effects of the drive field on resolution.

The lower drive field amplitude severely limited the ability to detect ProMag-labeled cell samples. As few as 5,000 cells were detected using drive field amplitudes of 20 x 20 mT. However, only 50,000 cells could be detected using drive field amplitudes of 5 x 5 mT. This was because the noise level was higher with the lower amplitude, reducing SNR. The value for 5*SD noise measured from the empty bed image acquired with drive field amplitudes of 5 x 5 mT was 5 times higher than for the empty bed image acquired with drive field amplitudes of 20 x 20 mT. For clinical scaling of MPI, peripheral nerve stimulation (PNS) is the main safety constraint of the drive field. PNS limits have been shown to decrease with frequency and has been observed in drive field amplitudes over 5 mT [77,78]. Our study has shown that a lower drive field greatly impedes the sensitivity and on a clinical system this loss in sensitivity will need to be recovered for cell tracking with optimal SPIO particles, efficient cell labeling, or longer scan times.

### *In Vivo* MPI Cell Detection Limits

The optimal parameters from the *in vitro* imaging of cell samples (3.0 T/m and 20 x 20 mT) were used to image mice injected with 2,000 or 5,000 ProMag-labeled cells. No signal was detected using 2D MPI in any of the mice. MPI signal was visible in the two mice injected with 5,000 cells using 3D MPI. The fact that we could detect 5,000 cells with 2D MPI *in vitro*, but not *in vivo*, could be related to the dispersion of cells from the injection site which reduces the cell density per voxel, leading some cells to fall below the detection limit. It could also be due to gut signal which could limit detection and quantification of lower signals due to the dynamic range limitations of MPI [20,47].

### Limitations

In this study, imaging of cell samples was only evaluated with single acquisition 2D MPI. Signal averaging is expected to improve sensitivity; however, this has not been well studied for MPI. Sehl *et al.* previously showed that in 2D MPI, a significant reduction in background noise was measured with 8 averages compared to 1 average, however, this did not improve cell detection [9]. 3D MPI using 35 projections did offer an improvement in sensitivity; 4,000 stem cells were detected with 2D MPI compared to 2,000 stem cells with 3D MPI. The acquisition time for 3D images (30 minutes) is approximately 15 times longer than for 2D images (2 minutes). For *in vivo* imaging, using 3D MPI to improve cell detection limits must be weighed against the amount of time that mice need to be anesthetized for a single scan, or the time required to image multiple mice in experimental groups. Our preliminary assessment of *in vivo* detection limits was only evaluated in a small number of mice and only immediately after the injection of cells. Future studies will include longitudinal imaging of mice to better understand how the MPI signal changes with the dispersion of cells *in vivo* or with cell death and the clearance of iron particles.

## Conclusion

In this study, we evaluated ProMag as a novel MPI cellular imaging tracer and compared it to two other commercial SPIO tracers that are more commonly used for MPI. We also studied the effect of changing imaging parameters on MPI sensitivity, resolution and cell detection limits with ProMag-labeled cells. For preclinical MPI cell tracking, ProMag, a polymer-encapsulated MPIO particle, has certain advantages including efficient cell labeling and high cellular iron loading. Furthermore, the MPI signal is unchanged after ProMag is compartmentalized in cells. All of these advantages lead to high MPI sensitivity for cell detection. Scanner-specific parameters can be adjusted for improved sensitivity or resolution when using ProMag, but careful consideration needs to be taken due to the trade-off between the two.

## Acknowledgements

We would like to acknowledge funding from the Canadian Institutes of Health Research, the National Sciences and Engineering Research Council of Canada, the (ImPaKT) Facility at Western University and the Canadian Foundation of Innovation. We would like to acknowledge Dr. Lauren Flynn for providing the cells used in this project and Dr. Olivia C. Sehl for her assistance in editing and reviewing this manuscript. We would also like to acknowledge the London Regional Flow Cytometry Facility for the use of their facility and support for the flow cytometry experiment in this study.

## References

1. Daley, G.Q. The Promise and Perils of Stem Cell Therapeutics. Cell Stem Cell 2012, 10, 740–749, doi:10.1016/j.stem.2012.05.010.

2. Squillaro, T.; Peluso, G.; Galderisi, U. Clinical Trials with Mesenchymal Stem Cells: An Update. Cell Transplant 2016, 25, 829–848, doi:10.3727/096368915X689622.

3. Han, J.; Zhang, B.; Zheng, S.; Jiang, Y.; Zhang, X.; Mao, K. The Progress and Prospects of Immune Cell Therapy for the Treatment of Cancer. Cell Transplant 2024, 33, doi:10.1177/09636897241231892.

4. Schots, R.; De Keulenaer, G.; Schoors, D.; Caveliers, V.; Dujardin, M.; Verheye, S.; Van Camp, G.; Franken, P.R.; Roland, J.; Van Riet, I.;, et al. Evidence That Intracoronary-Injected CD133+ Peripheral Blood Progenitor Cells Home to the Myocardium in Chronic Postinfarction Heart Failure. Exp Hematol 2007, 35, 1884–1890, doi:10.1016/j.exphem.2007.07.012.

5. Kurpisz, M.; Czepczyński, R.; Grygielska, B.; Majewski, M.; Fiszer, D.; Jerzykowska, O.; Sowiński, J.; Siminiak, T. Bone Marrow Stem Cell Imaging after Intracoronary Administration. Int J Cardiol 2007, 121, 194–195, doi:10.1016/j.ijcard.2006.08.062.

6. Han, Y.; Li, Y.; Wu, W.E.; Liu, Z. Noninvasive Strategies of Cell-Tracking in Vivo. TrAC Trends in Analytical Chemistry 2024, 172, 117616, doi:10.1016/j.trac.2024.117616.

7. Ahrens, E.T.; Bulte, J.W.M. Tracking Immune Cells in Vivo Using Magnetic Resonance Imaging. Nat Rev Immunol 2013, 13, 755–763, doi:10.1038/nri3531.

8. Gaudet, J.M.; Ribot, E.J.; Chen, Y.; Gilbert, K.M.; Foster, P.J. Tracking the Fate of Stem Cell Implants with Fluorine-19 MRI. PLoS One 2015, 10, e0118544, doi:10.1371/journal.pone.0118544.

9. Sehl, O.C.; Foster, P.J. The Sensitivity of Magnetic Particle Imaging and Fluorine-19 Magnetic Resonance Imaging for Cell Tracking. Sci Rep 2021, 11, 22198, doi:10.1038/s41598-021-01642-3.

10. Kiraga, Ł.; Kucharzewska, P.; Paisey, S.; Cheda, Ł.; Domańska, A.; Rogulski, Z.; Rygiel, T.P.; Boffi, A.; Król, M. Nuclear Imaging for Immune Cell Tracking in Vivo – Comparison of Various Cell Labeling Methods and Their Application. Coord Chem Rev 2021, 445, 214008, doi:10.1016/j.ccr.2021.214008.

11. Sehl, O.C.; Makela, A. V.; Hamilton, A.M.; Foster, P.J. Trimodal Cell Tracking In Vivo: Combining Iron- and Fluorine-Based Magnetic Resonance Imaging with Magnetic Particle Imaging to Monitor the Delivery of Mesenchymal Stem Cells and the Ensuing Inflammation. Tomography 2019, 5, 367–376, doi:10.18383/j.tom.2019.00020.

12. Bulte, J.W.M.; Walczak, P.; Janowski, M.; Krishnan, K.M.; Arami, H.; Halkola, A.; Gleich, B.; Rahmer, J. Quantitative “Hot-Spot” Imaging of Transplanted Stem Cells Using Superparamagnetic Tracers and Magnetic Particle Imaging. Tomography 2015, 1, 91–97, doi:10.18383/j.tom.2015.00172.

13. Zheng, B.; Vazin, T.; Goodwill, P.W.; Conway, A.; Verma, A.; Ulku Saritas, E.; Schaffer, D.; Conolly, S.M. Magnetic Particle Imaging Tracks the Long-Term Fate of in Vivo Neural Cell Implants with High Image Contrast. Sci Rep 2015, 5, 14055, doi:10.1038/srep14055.

14. Shalaby, N.; Kelly, J.J.; Sehl, O.C.; Gevaert, J.J.; Fox, M.S.; Qi, Q.; Foster, P.J.; Thiessen, J.D.; Hicks, J.W.; Scholl, T.J.;, et al. Complementary Early-Phase Magnetic Particle Imaging and Late-Phase Positron Emission Tomography Reporter Imaging of Mesenchymal Stem Cells *in Vivo*. Nanoscale 2023, 15, 3408–3418, doi:10.1039/D2NR03684C.

15. Fink, C.; Gevaert, J.J.; Barrett, J.W.; Dikeakos, J.D.; Foster, P.J.; Dekaban, G.A. In Vivo Tracking of Adenoviral-Transduced Iron Oxide-Labeled Bone Marrow-Derived Dendritic Cells Using Magnetic Particle Imaging. Eur Radiol Exp 2023, 7, 42, doi:10.1186/s41747-023-00359-4.

16. Mangarova, D.B.; Brangsch, J.; Mohtashamdolatshahi, A.; Kosch, O.; Paysen, H.; Wiekhorst, F.; Klopfleisch, R.; Buchholz, R.; Karst, U.; Taupitz, M.;, et al. Ex Vivo Magnetic Particle Imaging of Vascular Inflammation in Abdominal Aortic Aneurysm in a Murine Model. Sci Rep 2020, 10, 12410, doi:10.1038/s41598-020-69299-y.

17. Sehl, O.C.; Yang, Y.; Anjier, A.R.; Nevozhay, D.; Cheng, D.; Guo, K.; Fellows, B.; Mohtasebzadeh, A.R.; Mason, E.E.; Sanders, T.;, et al. Labeling Natural Killer Cells with Superparamagnetic Iron Oxide Nanoparticles for Detection by Preclinical and Clinical-Scale Magnetic Particle Imaging 2024.

18. Rivera-Rodriguez, A.; Hoang-Minh, L.B.; Chiu-Lam, A.; Sarna, N.; Marrero-Morales, L.; Mitchell, D.A.; Rinaldi-Ramos, C.M. Tracking Adoptive T Cell Immunotherapy Using Magnetic Particle Imaging. Nanotheranostics 2021, 5, 431–444, doi:10.7150/ntno.55165.

19. Kiru, L.; Zlitni, A.; Tousley, A.M.; Dalton, G.N.; Wu, W.; Lafortune, F.; Liu, A.; Cunanan, K.M.; Nejadnik, H.; Sulchek, T.;, et al. In Vivo Imaging of Nanoparticle-Labeled CAR T Cells. Proceedings of the National Academy of Sciences 2022, 119, doi:10.1073/pnas.2102363119.

20. Fernando, N.; Gevaert, J.J.; Konkle, J.; Goodwill, P.; Foster, P.J. Focused Small Field of View Magnetic Particle Imaging for the Isolation and Quantification of MPI Signal in Tumours. Int J Magn Part Imaging 2024, 10, 11, doi:10.18416/ijmpi.2024.2411001.

21. Williams, R.J.; Sehl, O.C.; Gevaert, J.J.; Liu, S.; Kelly, J.J.; Foster, P.J.; Ronald, J.A. Dual Magnetic Particle Imaging and Akaluc Bioluminescence Imaging for Tracking Cancer Cell Metastasis. Tomography 2023, 9, 178–194, doi:10.3390/tomography9010016.

22. Melo, K.P.; Makela, A. V.; Knier, N.N.; Hamilton, A.M.; Foster, P.J. Magnetic Microspheres Can Be Used for Magnetic Particle Imaging of Cancer Cells Arrested in the Mouse Brain. Magn Reson Med 2022, 87, 312–322, doi:10.1002/mrm.28987.

23. Makela, A. V.; Schott, M.A.; Sehl, O.C.; Gevaert, J.J.; Foster, P.J.; Contag, C.H. Tracking the Fates of Iron-Labeled Tumor Cells *in Vivo* Using Magnetic Particle Imaging. Nanoscale Adv 2022, 4, 3617–3623, doi:10.1039/D2NA00008C.

24. Parkins, K.M.; Melo, K.P.; Chen, Y.; Ronald, J.A.; Foster, P.J. Visualizing Tumour Self-Homing with Magnetic Particle Imaging. Nanoscale 2021, 13, 6016–6023, doi:10.1039/D0NR07983A.

25. Toomajian, V.A.; Tundo, A.; Ural, E.E.; Greeson, E.M.; Contag, C.H.; Makela, A. V. Magnetic Particle Imaging Reveals That Iron-Labeled Extracellular Vesicles Accumulate in Brains of Mice with Metastases. ACS Appl Mater Interfaces 2024, 16, 30860–30873, doi:10.1021/acsami.4c04920.

26. Jung, K.O.; Jo, H.; Yu, J.H.; Gambhir, S.S.; Pratx, G. Development and MPI Tracking of Novel Hypoxia-Targeted Theranostic Exosomes. Biomaterials 2018, 177, 139–148, doi:10.1016/j.biomaterials.2018.05.048.

27. Graeser, M.; Thieben, F.; Szwargulski, P.; Werner, F.; Gdaniec, N.; Boberg, M.; Griese, F.; Möddel, M.; Ludewig, P.; van de Ven, D.;, et al. Human-Sized Magnetic Particle Imaging for Brain Applications. Nat Commun 2019, 10, doi:10.1038/s41467-019-09704-x.

28. Vogel, P.; Rückert, M.A.; Greiner, C.; Günther, J.; Reichl, T.; Kampf, T.; Bley, T.A.; Behr, V.C.; Herz, S. IMPI: Portable Human-Sized Magnetic Particle Imaging Scanner for Real-Time Endovascular Interventions. Sci Rep 2023, 13, 10472, doi:10.1038/s41598-023-37351-2.

29. Mason, E.E.; Cooley, C.Z.; Cauley, S.F.; Griswold, M.A.; Conolly, S.M.; Wald, L.L. Design Analysis of an MPI Human Functional Brain Scanner. Int J Magn Part Imaging 2017, 3, doi:10.18416/ijmpi.2017.1703008.

30. Mattingly, E.; Sliwiak, M.; Mason, E.E.; Chacon-Caldera, J.; Barksdale, A.C.; Niebel, F.H.; Herb, K.; Graeser, M.; Wald, L.L. rValidation of a Magnetic Particle Imaging (MPI) System for Human Brain Imaging. Phys Med Biol 2024, doi:10.1088/1361-6560/ad9db0.

31. Borgert, J.; Gleich, B.; Buzug, T.M. Magnetic Particle Imaging. In *Springer Handbook of Medical Technology*; Springer Berlin Heidelberg: Berlin, Heidelberg, 2011; pp. 461–476.

32. Borgert, J.; Schmidt, J.D.; Schmale, I.; Bontus, C.; Gleich, B.; David, B.; Weizenecker, J.; Jockram, J.; Lauruschkat, C.; Mende, O.;, et al. Perspectives on Clinical Magnetic Particle Imaging. Biomedizinische Technik/Biomedical Engineering 2013, 58, doi:10.1515/bmt-2012-0064.

33. Knopp, T.; Gdaniec, N.; Möddel, M. Magnetic Particle Imaging: From Proof of Principle to Preclinical Applications. Phys Med Biol 2017, 62, R124–R178, doi:10.1088/1361-6560/aa6c99.

34. Neumann, A.; Gräfe, K.; von Gladiss, A.; Ahlborg, M.; Behrends, A.; Chen, X.; Schumacher, J.; Blancke Soares, Y.; Friedrich, T.; Wei, H.;, et al. Recent Developments in Magnetic Particle Imaging. J Magn Magn Mater 2022, 550, 169037, doi:10.1016/j.jmmm.2022.169037.

35. Irfan, M.; Dogan, N. Comprehensive Evaluation of Magnetic Particle Imaging (MPI) Scanners for Biomedical Applications. IEEE Access 2022, 10, 86718–86732, doi:10.1109/ACCESS.2022.3197586.

36. Harvell-Smith, S.; Tung, L.D.; Thanh, N.T.K. Magnetic Particle Imaging: Tracer Development and the Biomedical Applications of a Radiation-Free, Sensitive, and Quantitative Imaging Modality. Nanoscale 2022, 14, 3658–3697, doi:10.1039/D1NR05670K.

37. Sehl, O.C.; Tiret, B.; Berih, M.A.; Makela, A. V.; Goodwill, P.W.; Foster, P.J. MPI Region of Interest (ROI) Analysis and Quantification of Iron in Different Volumes. Int J Magn Part Imaging 2022, 8.

38. Croft, L.R.; Goodwill, P.W.; Konkle, J.J.; Arami, H.; Price, D.A.; Li, A.X.; Saritas, E.U.; Conolly, S.M. Low Drive Field Amplitude for Improved Image Resolution in Magnetic Particle Imaging. Med Phys 2015, 43, 424–435, doi:10.1118/1.4938097.

39. Goodwill, P.W.; Conolly, S.M. The X-Space Formulation of the Magnetic Particle Imaging Process: 1-D Signal, Resolution, Bandwidth, SNR, SAR, and Magnetostimulation. IEEE Trans Med Imaging 2010, 29, 1851–1859, doi:10.1109/TMI.2010.2052284.

40. Erbe, M. Field Free Line Magnetic Particle Imaging; Springer Fachmedien Wiesbaden: Wiesbaden, 2014; ISBN 978-3-658-05336-9.

41. Tay, Z.W.; Hensley, D.W.; Chandrasekharan, P.; Zheng, B.; Conolly, S.M. Optimization of Drive Parameters for Resolution, Sensitivity and Safety in Magnetic Particle Imaging. IEEE Trans Med Imaging 2020, 39, 1724–1734, doi:10.1109/TMI.2019.2957041.

42. Weizenecker, J.; Borgert, J.; Gleich, B. A Simulation Study on the Resolution and Sensitivity of Magnetic Particle Imaging. Phys Med Biol 2007, 52, 6363–6374, doi:10.1088/0031-9155/52/21/001.

43. Liu, S.; Chiu-Lam, A.; Rivera-Rodriguez, A.; DeGroff, R.; Savliwala, S.; Sarna, N.; Rinaldi-Ramos, C.M. Long Circulating Tracer Tailored for Magnetic Particle Imaging. Nanotheranostics 2021, 5, 348–361, doi:10.7150/ntno.58548.

44. Mohtashamdolatshahi, A.; Kratz, H.; Kosch, O.; Hauptmann, R.; Stolzenburg, N.; Wiekhorst, F.; Sack, I.; Hamm, B.; Taupitz, M.; Schnorr, J. In Vivo Magnetic Particle Imaging: Angiography of Inferior Vena Cava and Aorta in Rats Using Newly Developed Multicore Particles. Sci Rep 2020, 10, 17247, doi:10.1038/s41598-020-74151-4.

45. Song, G.; Chen, M.; Zhang, Y.; Cui, L.; Qu, H.; Zheng, X.; Wintermark, M.; Liu, Z.; Rao, J. Janus Iron Oxides @ Semiconducting Polymer Nanoparticle Tracer for Cell Tracking by Magnetic Particle Imaging. Nano Lett 2018, 18, 182–189, doi:10.1021/acs.nanolett.7b03829.

46. Vogel, P.; Kampf, T.; Rückert, M.A.; Grüttner, C.; Kowalski, A.; Teller, H.; Behr, V.C. Synomag®: The New High-Performance Tracer for Magnetic Particle Imaging. Int J Magn Part Imaging 2021, 7, doi:10.18416/IJMPI.2021.2103003.

47. Gevaert, J.J.; Fink, C.; Dikeakos, J.D.; Dekaban, G.A.; Foster, P.J. Magnetic Particle Imaging Is a Sensitive In Vivo Imaging Modality for the Detection of Dendritic Cell Migration. Mol Imaging Biol 2022, doi:10.1007/s11307-022-01738-w.

48. Tay, Z.W.; Hensley, D.W.; Vreeland, E.C.; Zheng, B.; Conolly, S.M. The Relaxation Wall: Experimental Limits to Improving MPI Spatial Resolution by Increasing Nanoparticle Core Size. Biomed Phys Eng Express 2017, 3, 035003, doi:10.1088/2057-1976/aa6ab6.

49. Ferguson, R.M.; Khandhar, A.P.; Kemp, S.J.; Arami, H.; Saritas, E.U.; Croft, L.R.; Konkle, J.; Goodwill, P.W.; Halkola, A.; Rahmer, J.;, et al. Magnetic Particle Imaging With Tailored Iron Oxide Nanoparticle Tracers. IEEE Trans Med Imaging 2015, 34, 1077–1084, doi:10.1109/TMI.2014.2375065.

50. Eberbeck, D.; Wiekhorst, F.; Wagner, S.; Trahms, L. How the Size Distribution of Magnetic Nanoparticles Determines Their Magnetic Particle Imaging Performance. Appl Phys Lett 2011, 98, doi:10.1063/1.3586776.

51. Kahmann, T.; Ludwig, F. Magnetic Field Dependence of the Effective Magnetic Moment of Multi-Core Nanoparticles. J Appl Phys 2020, 127, doi:10.1063/5.0011629.

52. Gehrke, N.; Heinke, D.; Eberbeck, D.; Ludwig, F.; Wawrzik, T.; Kuhlmann, C.; Briel, A. Magnetic Characterization of Clustered Core Magnetic Nanoparticles for MPI. IEEE Trans Magn 2015, 51, 1–4, doi:10.1109/TMAG.2014.2358275.

53. Trozzo, S.; Neupane, B.; Foster, P.J. A Comparison of the Sensitivity and Cellular Detection Capabilities of Magnetic Particle Imaging and Bioluminescence Imaging. Tomography 2024, 10, 1846–1865, doi:10.3390/tomography10110135.

54. Calvert, N.D.; Yu, L.; Sehl, O.C.; Gevaert, J.J.; Knier, N.N.; Rivera-Rodriguez, A.; Goulet, C.S.; Fernando, N.; Flood, S.; Rinaldi-Ramos, C.M.;, et al. The Careful Selection of Zwitterionic Nanoparticle Coating Results in Rapid and Efficient Cell Labeling for Imaging-based Cell Tracking. Aggregate 2024, doi:10.1002/agt2.609.

55. Suzuka, H.; Mimura, A.; Inaoka, Y.; Murase, K. Magnetic Nanoparticles in Macrophages and Cancer Cells Exhibit Different Signal Behavior on Magnetic Particle Imaging. J Nanosci Nanotechnol 2019, 19, 6857–6865, doi:10.1166/jnn.2019.16619.

56. Arami, H.; Ferguson, R.M.; Khandhar, A.P.; Krishnan, K.M. Size-dependent Ferrohydrodynamic Relaxometry of Magnetic Particle Imaging Tracers in Different Environments. Med Phys 2013, 40, doi:10.1118/1.4810962.

57. Heyn, C.; Ronald, J.A.; Ramadan, S.S.; Snir, J.A.; Barry, A.M.; MacKenzie, L.T.; Mikulis, D.J.; Palmieri, D.; Bronder, J.L.; Steeg, P.S.;, et al. In Vivo MRI of Cancer Cell Fate at the Single-cell Level in a Mouse Model of Breast Cancer Metastasis to the Brain. Magn Reson Med 2006, 56, 1001–1010, doi:10.1002/mrm.21029.

58. Shapiro, E.M.; Skrtic, S.; Sharer, K.; Hill, J.M.; Dunbar, C.E.; Koretsky, A.P. MRI Detection of Single Particles for Cellular Imaging. Proceedings of the National Academy of Sciences 2004, 101, 10901–10906, doi:10.1073/pnas.0403918101.

59. Paysen, H.; Loewa, N.; Stach, A.; Wells, J.; Kosch, O.; Twamley, S.; Makowski, M.R.; Schaeffter, T.; Ludwig, A.; Wiekhorst, F. Cellular Uptake of Magnetic Nanoparticles Imaged and Quantified by Magnetic Particle Imaging. Sci Rep 2020, 10, 1922, doi:10.1038/s41598-020-58853-3.

60. Gevaert, J.J.; Kyle, ·; Beek, V.; Sehl, O.C.; Foster, P.J. VivoTrax+ ^TM^ Improves the Detection of Cancer Cells with Magnetic Particle Imaging. Int J Magn Part Imaging 2022, 8, 10, doi:10.18416/ijmpi.2022.2210001.

61. Arami, H.; Krishnan, K.M. Intracellular Performance of Tailored Nanoparticle Tracers in Magnetic Particle Imaging. J Appl Phys 2014, 115, doi:10.1063/1.4867756.

62. Them, K.; Salamon, J.; Szwargulski, P.; Sequeira, S.; Kaul, M.G.; Lange, C.; Ittrich, H.; Knopp, T. Increasing the Sensitivity for Stem Cell Monitoring in System-Function Based Magnetic Particle Imaging. Phys Med Biol 2016, 61, 3279–3290, doi:10.1088/0031-9155/61/9/3279.

63. Sehl, O.C.; Gevaert, J.J.; Melo, K.P.; Knier, N.N.; Foster, P.J. A Perspective on Cell Tracking with Magnetic Particle Imaging. Tomography 2020, 6, 315–324, doi:10.18383/j.tom.2020.00043.

64. Thu, M.S.; Bryant, L.H.; Coppola, T.; Jordan, E.K.; Budde, M.D.; Lewis, B.K.; Chaudhry, A.; Ren, J.; Varma, N.R.S.; Arbab, A.S.;, et al. Self-Assembling Nanocomplexes by Combining Ferumoxytol, Heparin and Protamine for Cell Tracking by Magnetic Resonance Imaging. Nat Med 2012, 18, 463–467, doi:10.1038/nm.2666.

65. Burgess, A.E. The Rose Model, Revisited. Journal of the Optical Society of America A 1999, 16, 633, doi:10.1364/JOSAA.16.000633.

66. Rose, A. The Sensitivity Performance of the Human Eye on an Absolute Scale*. J Opt Soc Am 1948, 38, 196, doi:10.1364/JOSA.38.000196.

67. Guzy, J.; Chakravarty, S.; Buchanan, F.J.; Chen, H.; Gaudet, J.M.; Hix, J.M.L.; Mallett, C.L.; Shapiro, E.M. Complex Relationship between Iron Oxide Nanoparticle Degradation and the Signal Intensity in Magnetic Particle Imaging. ACS Appl Nano Mater 2020, 3, 3991–3999, doi:10.1021/acsanm.0c00779.

68. Poller, W.C.; Löwa, N.; Wiekhorst, F.; Taupitz, M.; Wagner, S.; Möller, K.; Baumann, G.; Stangl, V.; Trahms, L.; Ludwig, A. Magnetic Particle Spectroscopy Reveals Dynamic Changes in the Magnetic Behavior of Very Small Superparamagnetic Iron Oxide Nanoparticles During Cellular Uptake and Enables Determination of Cell-Labeling Efficacy. J Biomed Nanotechnol 2016, 12, 337–346, doi:10.1166/jbn.2016.2204.

69. Teeman, E.; Shasha, C.; Evans, J.E.; Krishnan, K.M. Intracellular Dynamics of Superparamagnetic Iron Oxide Nanoparticles for Magnetic Particle Imaging. Nanoscale 2019, 11, 7771–7780, doi:10.1039/C9NR01395D.

70. Tang, K.S.; Hann, B.; Shapiro, E.M. On the Use of Micron-Sized Iron Oxide Particles (MPIOS) to Label Resting Monocytes in Bone Marrow. Mol Imaging Biol 2011, 13, 819– 824, doi:10.1007/s11307-010-0437-3.

71. Boulland, J.-L.; Leung, D.S.Y.; Thuen, M.; Vik-Mo, E.; Joel, M.; Perreault, M.-C.; Langmoen, I.A.; Haraldseth, O.; Glover, J.C. Evaluation of Intracellular Labeling with Micron-Sized Particles of Iron Oxide (MPIOs) as a General Tool for In Vitro and in Vivo Tracking of Human Stem and Progenitor Cells. Cell Transplant 2012, 21, 1743–1759, doi:10.3727/096368911X627598.

72. Rohani, R.; de Chickera, S.N.; Willert, C.; Chen, Y.; Dekaban, G.A.; Foster, P.J. In Vivo Cellular MRI of Dendritic Cell Migration Using Micrometer-Sized Iron Oxide (MPIO) Particles. Mol Imaging Biol 2011, 13, 679–694, doi:10.1007/s11307-010-0403-0.

73. Chaumeil, M.M.; Gini, B.; Yang, H.; Iwanami, A.; Sukumar, S.; Ozawa, T.; Pieper, R.O.; Mischel, P.S.; James, C.D.; Berger, M.S.;, et al. Longitudinal Evaluation of MPIO-Labeled Stem Cell Biodistribution in Glioblastoma Using High Resolution and Contrast-Enhanced MR Imaging at 14.1Tesla. Neuro Oncol 2012, 14, 1050–1061, doi:10.1093/neuonc/nos126.

74. Shapiro, E.M. Biodegradable, Polymer Encapsulated, Metal Oxide Particles for MRI-based Cell Tracking. Magn Reson Med 2015, 73, 376–389, doi:10.1002/mrm.25263.

75. Perez-Balderas, F.; van Kasteren, S.I.; Aljabali, A.A.A.; Wals, K.; Serres, S.; Jefferson, A.; Sarmiento Soto, M.; Khrapitchev, A.A.; Larkin, J.R.; Bristow, C.;, et al. Covalent Assembly of Nanoparticles as a Peptidase-Degradable Platform for Molecular MRI. Nat Commun 2017, 8, 14254, doi:10.1038/ncomms14254.

76. 76. U.S. Department of Health and Human Services; Food and Drug Administration. (2008). Guidance for FDA Reviewers and Sponsors: Content and Review of Chemistry, manufacturing, and Control (CMC) Information for Human Somatic Cell Therapy Investigational New Drug Applications (INDs). https://www.fda.gov/regulatory-information/search-fda-guidance-documents/content-and-review-chemistry-manufacturing-and-control-cmc-information-human-somatic-cell-therapy

77. Barksdale, A.; Ferris, N.; Mattingly, E.; Sliwiak, M.; Guerin, B.; Wald, L.; Davids, M.; Klein, V. Measured PNS Thresholds in a Human Head MPI Solenoid from 200 Hz to 88.1 KHz 2024, 10.

78. Barksdale, A.C.; Ferris, N.G.; Mattingly, E.; Śliwiak, M.; Guerin, B.; Wald, L.L.; Davids, M.; Klein, V. Measurement of Peripheral Nerve Magnetostimulation Thresholds of a Head Solenoid Coil between 200 Hz and 88.1 KHz 2024.

